# What an atlas-fitted connectome can and cannot do: evolutionary search over interneuron stimulation in a whole-body *C. elegans* model

**DOI:** 10.64898/2026.09.06.749731

**Authors:** Taehee Lee

**Affiliations:** Department of Integrative Medicine, Yonsei University, Seoul, Republic of Korea

## Abstract

The *C. elegans* connectome is complete, but which behaviors its wiring can produce, and which it cannot, is unknown. We asked this question in a whole-body simulation whose only adaptive element is a learning agent outside the nervous system. The agent may inject current into a short whitelist of interneurons, never into motor neurons or muscles; the 302-neuron OpenWorm c302 network, with dynamics fitted once to a signal-propagation atlas, a literature-based motor layer and a viscoelastic body on agar do the rest. Every hypothesis was pre-registered with its judgment criterion. Stimulating five command and steering interneurons (AVB, AVA, SMDD, SMDV, RIV) was sufficient for chemotaxis to a source 2.5 mm away (91% of trials; four of five training seeds), whereas stimulating eight sensory neurons was no better than chance, reproducing at the behavioral level the silence of the sensory-to-command step in the fitted network. Evolution rediscovered the pirouette rule, reversing and executing a deep bend when concentration fell, and ablations showed that reversal, deep bend and steering were all necessary. Shuffling the wiring while preserving weights, counts and signs cut performance to a fifth; in the atlas-fitted dynamics this loss was carried by the gap-junction layer. In corridors four body widths wide, corners were passed by a deep bend pressed against the wall, so the dorsal or ventral direction of the bend fixed the direction of the turn. With ventral bends only, right turns almost never occurred and the worm entered the ventral arm of a T-maze first. Because temporal sensing cannot distinguish the arms at a junction, the learned strategy was to enter one arm and reverse when the odor faded; a left–right concentration difference was needed before the bend direction followed the source. The model yields three testable predictions for narrow-maze assays and separates what the wiring contributes to navigation from what sensing and the body must supply.

**Author summary:** Knowing every connection in a nervous system does not tell us what the wiring does. We took the complete wiring diagram of the worm *C. elegans*, kept every connection fixed, and let a learning algorithm discover what it could achieve by stimulating a handful of neurons, as an experimenter with optogenetic tools could. The algorithm was forbidden to touch the neurons that drive muscles. Through the wiring it learned to find an odor source, and it did so by rediscovering the strategy real worms use: reverse and turn sharply when the smell gets weaker. It failed when given only sensory neurons, which tells us that the sensory-to-command step is missing from the fitted wiring. Randomizing the wiring destroyed the behavior, and in our fitted model the electrical connections were the ones that mattered. In narrow mazes, the worm could only turn toward the side of its sharp bend, so a worm that bends to one side almost always turns that way and needs a costly reversal to reach food on the other side. These results separate what the wiring contributes from what the body and the senses contribute, and they make predictions that can be tested with worms in mazes.

## Introduction

The wiring diagram of *C. elegans* has been complete for decades [1, 2], but a wiring diagram is not a functional model, and the electrophysiological parameters that would turn one into the other are largely unknown [3, 4]. Two responses to this gap dominate. One fits network dynamics to measured signal propagation; the recent atlas of Randi et al. showed that anatomy alone predicts only part of the functional map [5]. The other fills the unknown parameters by optimization against behavior and obtains an ensemble of models consistent with anatomy [3]. Both ask what the wiring *does*. Here we ask a complementary question: with the wiring fixed and the free parameters confined to a learning agent outside the nervous system, which behaviors can be produced through the connectome, and which cannot?

We built a whole-body simulator in which the OpenWorm c302 network [4] is coupled to the neuromechanical motor layer of Boyle, Berri and Cohen [6] and to a two-dimensional viscoelastic body crawling on agar, with walls. The network dynamics were fitted once to the signal-propagation atlas [5] and then frozen. A small policy, trained by evolution strategies [7], observes the odor at the nose and may inject current into a whitelist of sensory, command and steering interneurons. It may never stimulate motor neurons or muscles. Behaviour therefore arises only from what the stimulated interneurons can do through the fixed wiring, the situation of an optogenetic experiment on a circuit whose downstream connectivity cannot be changed.

This framing turns the fitted connectome into an experimental subject. Because the agent adapts and the model does not, a failure to learn a behavior is evidence about the fitted network rather than about a hand-tuned controller, and a success identifies which interneurons are sufficient. It also lets us compare the learned strategy with those of real animals. Chemotaxis in *C. elegans* is produced by pirouettes, sharp reorientations whose probability rises when the concentration falls [8], and by a weathervane mechanism that steers gradually toward higher concentration [9]. Sharp reorientations are executed by a stereotyped neural sequence in which SMD and RIV drive the omega bend and SMDD versus SMDV set its dorsal or ventral direction, which the animal chooses according to the odor gradient [10]. Real animals in narrow T-mazes show no innate side bias and find a food arm in 85% of trials, a behavior that requires mechanosensation [11]. These anchors let us judge whether an agent constrained by the connectome converges on the same solutions.

We pre-registered each hypothesis with its judgment criterion before training and kept every negative result. We report four results. Five command and steering interneurons suffice for chemotaxis while sensory neurons do not. Evolution rediscovers the pirouette rule, and reversal, deep bend and steering are all required. In narrow corridors the direction of the deep bend is the direction of the turn, which makes arm choice at a T-junction a problem of bend direction and lateral sensing rather than of the connectome. The real wiring outperforms shuffled wiring through its gap-junction layer.

## Results

### The agent”s only lever is current into whitelisted interneurons

Everything downstream of the stimulated neurons is fixed (Fig 1). Every 0.5 s a small neural-network policy maps the recent history of log odor concentration at the nose, and in maze tasks the wall contact of the head, to currents of 0–6 pA injected into the chosen neurons. The network is the c302 connectome (302 neurons, 2,279 neuron-to-neuron chemical synapses, 1,084 gap junctions) with dynamics fitted to the signal-propagation atlas [5] (“Vfit”, Methods). The motor layer reads only five interneurons. Forward and backward locomotion are gated by AVB and AVA depolarization [6]; the SMDD − SMDV difference steers the head; and a RIV depolarization above threshold fires a traveling curvature pulse that produces an omega turn of about 170°, which we call the deep bend [10]. Under the rule adopted for maze experiments, the pulse is dorsal when SMDD exceeds SMDV and ventral otherwise, as in the neural sequence of directed turning [10]; the earlier ventral-only rule serves as a control. The two rules are recorded as architecture decision records ADR-016 and ADR-017 (Methods). Policies were trained by evolution strategies with a near-source curriculum and evaluated on 128–256 fresh random starts. Hypothesis identifiers (H1, H5-3, …) refer to Table 1.

**Table 1:**
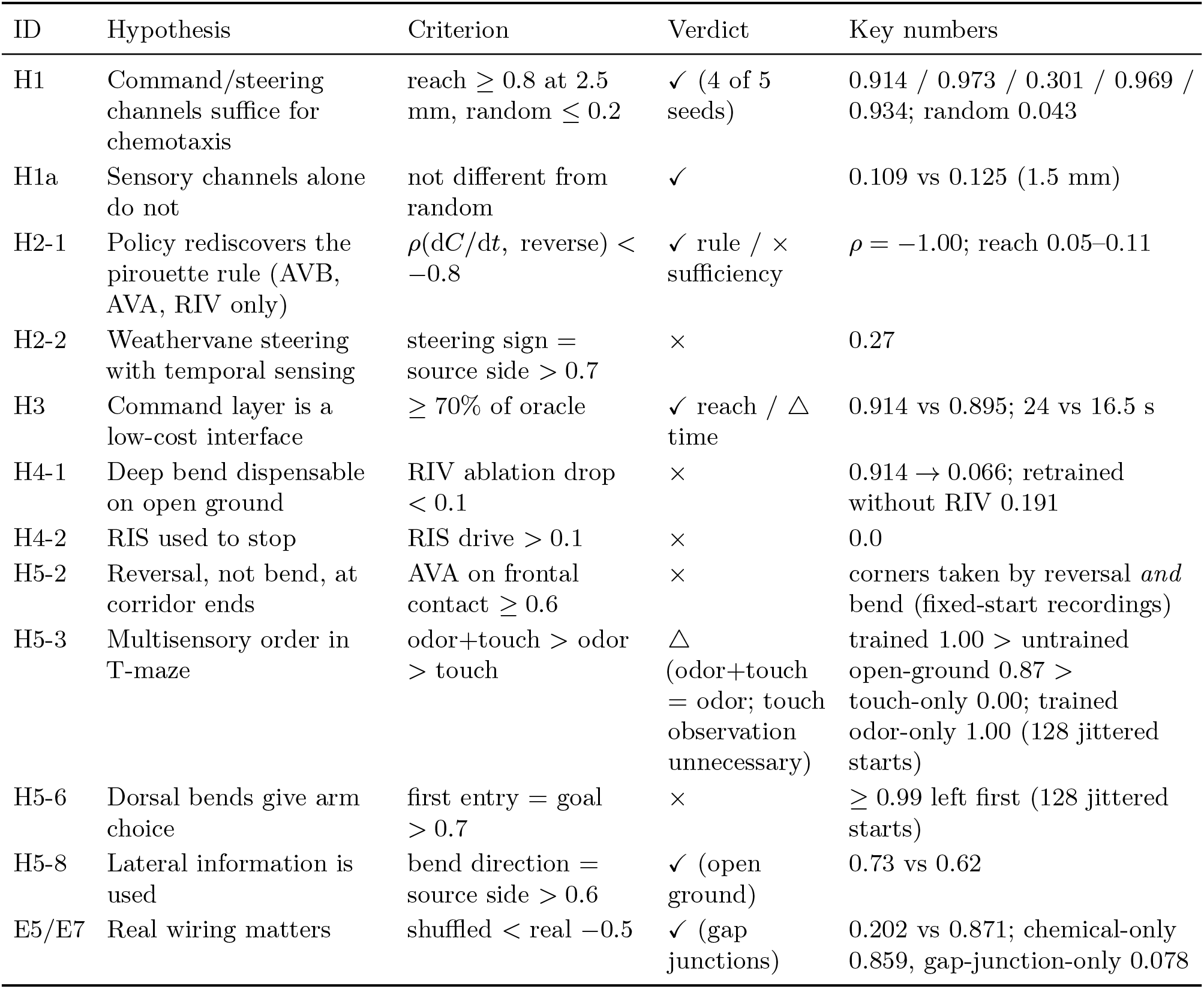
Pre-registered hypotheses and verdicts. *n* = 256 episodes unless stated.

| ID | Hypothesis | Criterion | Verdict | Key numbers |
| --- | --- | --- | --- | --- |
| H1 | Command/steering channels suffice for chemotaxis | reach $\geq 0.8$ at 2.5 mm, random $\leq 0.2$ | ✓ (4 of 5 seeds) | 0.914 / 0.973 / 0.301 / 0.969 / 0.934; random 0.043 |
| H1a | Sensory channels alone do not | not different from random | ✓ | 0.109 vs 0.125 (1.5 mm) |
| H2-1 | Policy rediscovers the pirouette rule (AVB, AVA, RIV only) | $\rho(dC/dt, \text{reverse}) < -0.8$ | ✓ rule / × sufficiency | $\rho = -1.00$; reach 0.05–0.11 |
| H2-2 | Weathervane steering with temporal sensing | steering sign = source side $> 0.7$ | × | 0.27 |
| H3 | Command layer is a low-cost interface | $\geq 70\%$ of oracle | ✓ reach / $\Delta$ time | 0.914 vs 0.895; 24 vs 16.5 s |
| H4-1 | Deep bend dispensable on open ground | RIV ablation drop $< 0.1$ | × | 0.914 → 0.066; retrained without RIV 0.191 |
| H4-2 | RIS used to stop | RIS drive $> 0.1$ | × | 0.0 |
| H5-2 | Reversal, not bend, at corridor ends | AVA on frontal contact $\geq 0.6$ | × | corners taken by reversal *and* bend (fixed-start recordings) |
| H5-3 | Multisensory order in T-maze | odor+touch $>$ odor $>$ touch | $\Delta$ (odor+touch = odor; touch observation unnecessary) | trained 1.00 $>$ untrained open-ground 0.87 $>$ touch-only 0.00; trained odor-only 1.00 (128 jittered starts) |
| H5-6 | Dorsal bends give arm choice | first entry = goal $> 0.7$ | × | $\geq 0.99$ left first (128 jittered starts) |
| H5-8 | Lateral information is used | bend direction = source side $> 0.6$ | ✓ (open ground) | 0.73 vs 0.62 |
| E5/E7 | Real wiring matters | shuffled $<$ real $-0.5$ | ✓ (gap junctions) | 0.202 vs 0.871; chemical-only 0.859, gap-junction-only 0.078 |

**Figure 1:**
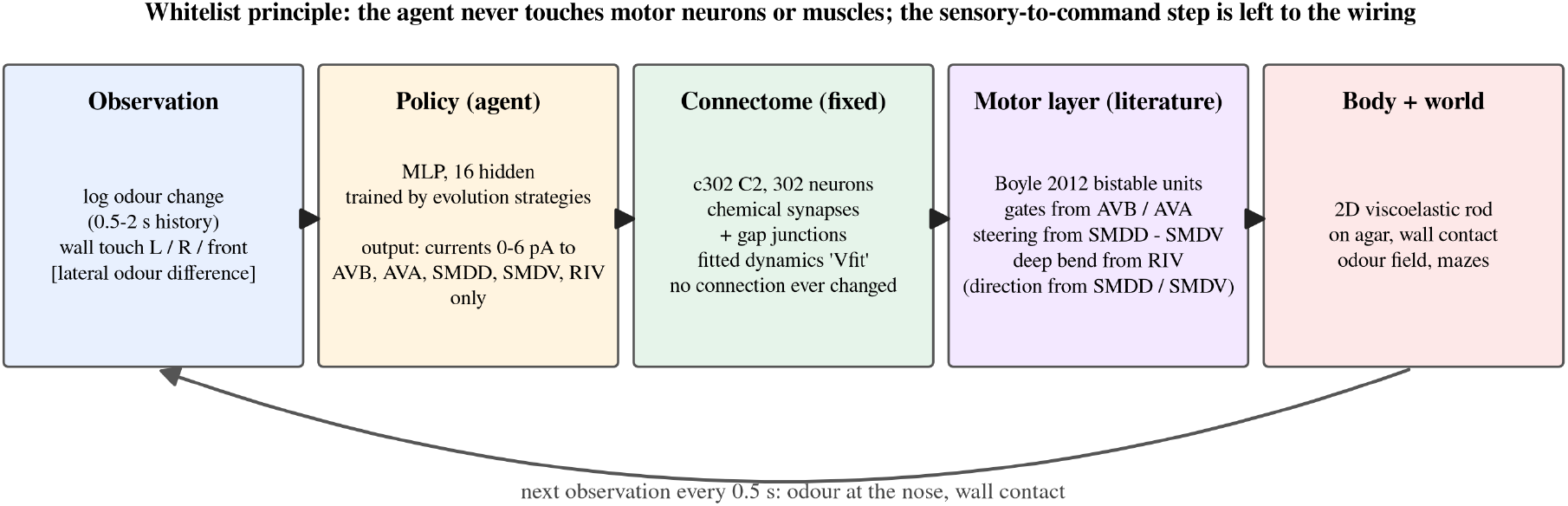
The pipeline and the whitelist principle. Observation → policy → stimulation of whitelisted interneurons → fixed c302 connectome with fitted dynamics → literature motor layer → viscoelastic body in an odor field with walls.

### Five command and steering interneurons suffice for chemotaxis; sensory neurons do not

Stimulating only AVB, AVA, SMDD, SMDV and RIV was sufficient for chemotaxis (Fig 2A). The trained policy reached a source 2.5 mm away within 40 s in 91.4% of 256 trials (Wilson 95% CI 0.873–0.943; median 24 s), against 4.3% for a random policy and 0% for a still worm; at 1.5 mm it reached 99.6%. Four of five independent training seeds cleared the pre-registered criterion of 0.8 (0.914, 0.973, 0.969 and 0.934; mean of five 0.82 *±* 0.29 s.d.). The remaining seed converged to a strategy that reversed on decline but never used the deep bend, reached 0.301, and stayed there after 100 further generations (0.324); we return to this local optimum below. An oracle controller that knew the source direction and steered the same body directly reached the source in 89.5% of trials, fewer than the learned policy, though faster (median 16.5 s versus 24 s). The oracle is a hand-tuned baseline, so we conclude only that the interneuron interface costs nothing in reach rate, not that it is optimal. Learning failed without three ingredients that we record as negative results: a living cost and a terminal distance penalty (otherwise standing still is a local optimum), relative rather than absolute concentration observations, and the curriculum (direct training at 2.5 mm reached 8% after 45 generations).

**Figure 2:**
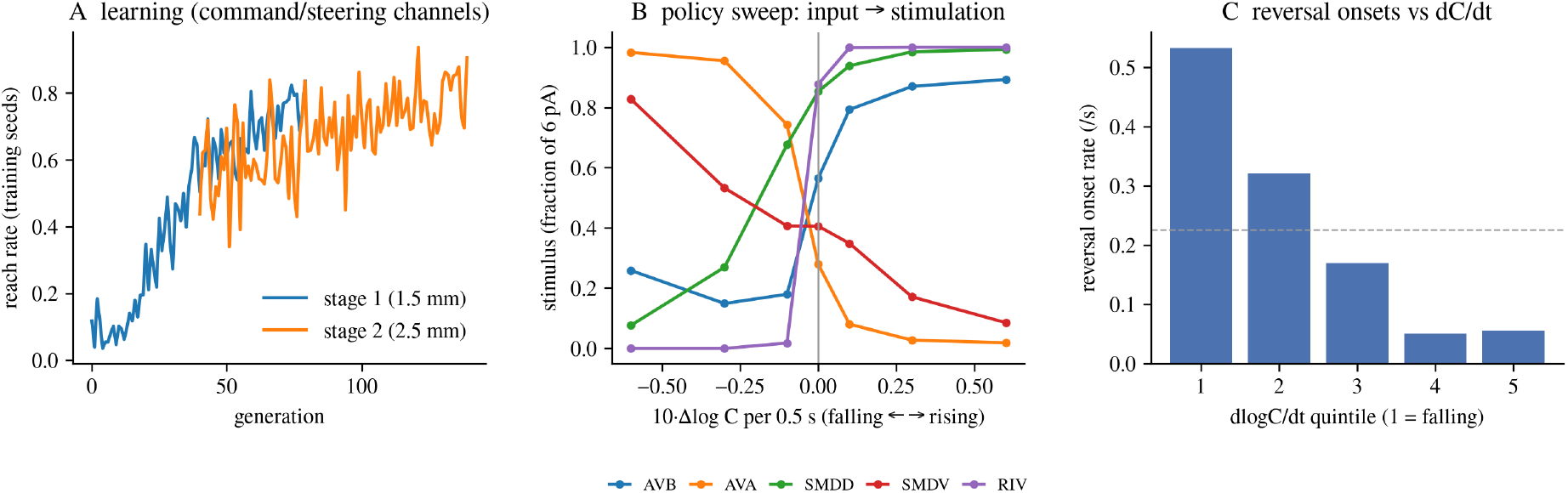
Chemotaxis through command and steering interneurons. (A) Reach rate of the training seeds over generations under the curriculum (stage 1 at 1.5 mm, stage 2 at 2.5 mm). (B) Policy sweep: stimulation of each channel as a function of the observed concentration change. (C) Reversal-onset rate by d*C/*d*t* quintile (1 = falling); the dashed line is the overall rate. Compare Pierce-Shimomura et al. [8].

Stimulating sensory neurons instead of interneurons did not produce chemotaxis. With ASE, AWC, AWA, ASK, ASH, PLM, ALM and AVM as the only channels, the best policy reached a source 1.5 mm away in 10.9% of trials (CI 0.077–0.154), against 12.5% for a random policy (CI 0.090–0.171), and its learning curve stayed at the reward of a still worm for 80 generations. The policy had tried to exploit a sub-millivolt asymmetry of the touch circuit, with the sign opposite to the biological one. This behavioral failure matches two properties of the fitted network established when it was built (Methods): static sensory stimulation does not differentiate the forward and backward command states, and 83% of the inhibitory responses in the atlas connect neuron pairs that have no one- or two-step synaptic path. The gap between the two conditions at 1.5 mm, 0.996 versus 0.109, is the contribution of the single missing step, sensory-to-command selection.

### Evolution rediscovers the pirouette rule, and reversal, deep bend and steering are all necessary

The learned policy implemented the pirouette rule of Pierce-Shimomura et al. [8] (Fig 2B–C). Sweeping the policy over the observed concentration change gave a clean switch. When concentration fell, AVA (reverse) was driven at 0.98 of maximum with a ventral head bias (SMDV 0.83); when it rose, AVB (forward), RIV (deep bend) and SMDD were driven at 0.89, 1.0 and 0.99. The rate of reversal onsets fell monotonically across quintiles of d*C/*d*t*, from 0.533 to 0.056 per second (ratio ≈ 9, Spearman *ρ* = −0.90), the sign and monotonicity reported for real animals [8]. The model”s baseline rate (0.23 per second) is about eight times the real one (0.023–0.030 per second [8]) because its reversal bouts are short (2 s) and frequent. The other successful seeds implemented the same switch with different weights. In all four navigating seeds the realized probability of the backward gate was highest in the falling quintile and lowest in the rising one (0.15 → 0.09, 0.30 → 0.04, 0.50 → 0.02 and 0.31 → 0.03). The onset-rate statistic, which counts AVA crossings of half maximum, was monotone in only two of them (*ρ* = −0.90 and −0.70; −0.30 and +0.70 in the others). The seed with the positive value reversed by switching AVB off while holding AVA near half maximum, so that the backward gate won without an AVA crossing. The rule is therefore reproduced at the level of gating in every navigating seed, and the crossing-based onset rate is a readout that depends on how the policy implements it. The weathervane mechanism [9] did not appear: the sign of head steering agreed with the side of the source in 0.27 of steps (criterion *>* 0.7), because steering was tied to d*C/*d*t*, not to the gradient direction, which a nose sampling only the temporal history cannot observe.

The rule alone was not sufficient. Restricting the channels to AVB, AVA and RIV, the pre-registered condition for the rule test, produced the rule in its purest form, reverse plus deep bend on decline and forward run on rise (*ρ* = −1.00), in two of three seeds, yet these policies could not navigate (reach 0.05–0.11 at 2.5 mm). Without steering the only reorientation is a fixed bend of about 170°, so the worm retraces its path. The third seed of this rule test found an alternative, a bend held continuously during runs, that reached 0.328.

All three components were necessary (Fig 3). Masking one channel of the trained policy at evaluation reduced the reach rate from 0.914 to 0.098 (AVA), 0.066 (RIV), 0.273 (SMDD and SMDV together), 0.434 (AVB) and 0.410 or 0.621 (SMDD or SMDV alone). Retraining without RIV reached 0.191 and retraining without steering 0.05–0.33 across three seeds (Fig 3B shows the best), so the necessity is not an artifact of the particular policy. Our pre-registered prediction that the deep bend would be dispensable on open ground was therefore false: with temporal sensing only, chemotaxis by modulating the rate of reorientation (klinokinesis) requires a large reorientation. A further pre-registered hypothesis asked whether the sleep-active neuron RIS [12] would be recruited to stop at the source. It was not. When the task rewarded dwelling at the source, with or without a movement cost, the policy stopped by withdrawing command drive below the gate threshold and RIS stimulation stayed at zero; RIS can only matter when the forward drive is not under the agent”s control.

**Figure 3:**
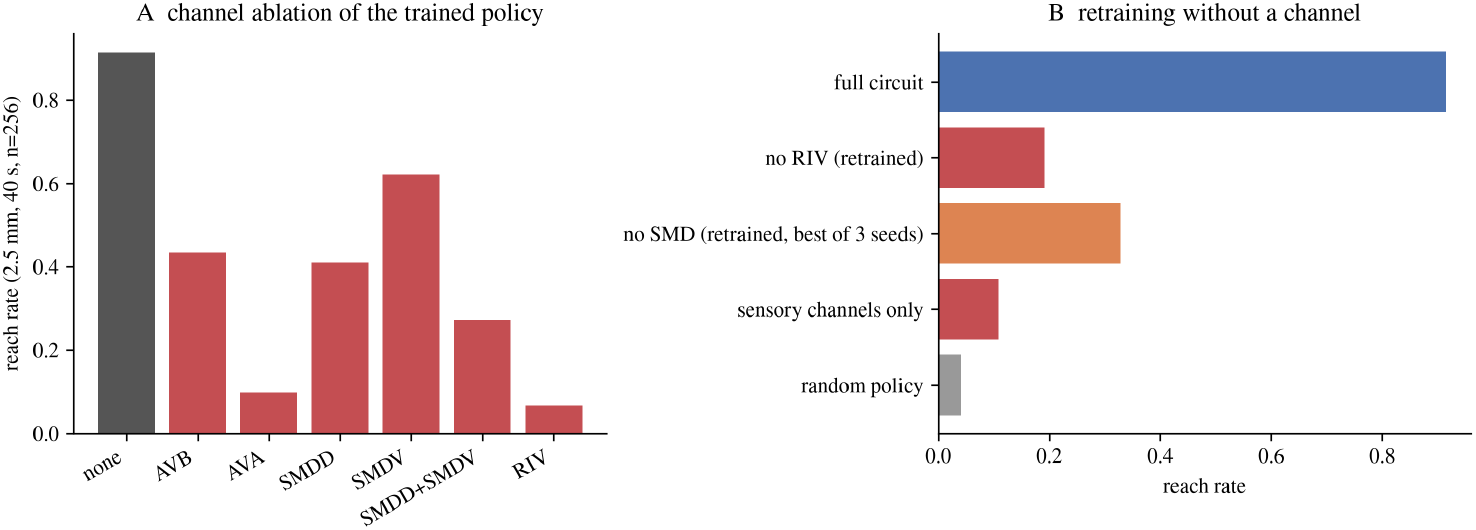
The minimal circuit. (A) Reach rate at 2.5 mm (*n* = 256) after masking each channel of the trained policy. (B) Reach rate after retraining without RIV or without SMD steering (best of three seeds), with the sensory-only and random policies for comparison.

### In narrow corridors the direction of the deep bend is the direction of the turn

In every recording, corners were passed by one mechanism: a deep bend pressed against the wall (Fig 4). We added a wall potential to the body model and lattice mazes with corridors 0.3 mm wide, about four body widths. In corridors of this width the omega bend, whose free diameter is 0.34 mm, is blocked (reorientation +2° instead of +170°) and forward speed halves (0.07–0.10 versus 0.15–0.18 mm/s). Yet the odor policy trained on open ground, placed in an L-shaped corridor without further training, passed the corner in 97–100% of 32 jittered starts at each width (Methods). Frame-by-frame traces showed the sequence stall → reversal → forward run with a deep-bend pulse pressed against the wall, after which the body slides around the corner. In the fixed-start recordings of the open-ground policy and of a maze-trained policy, a reversal command and a bend pulse both occurred within the 3 s before corner passage. Head steering alone could not turn the corner: in fixed-start tests, constant SMDD or SMDV bias with forward drive never passed the corner, and masking RIV reduced T-maze completion to 0.

**Figure 4:**
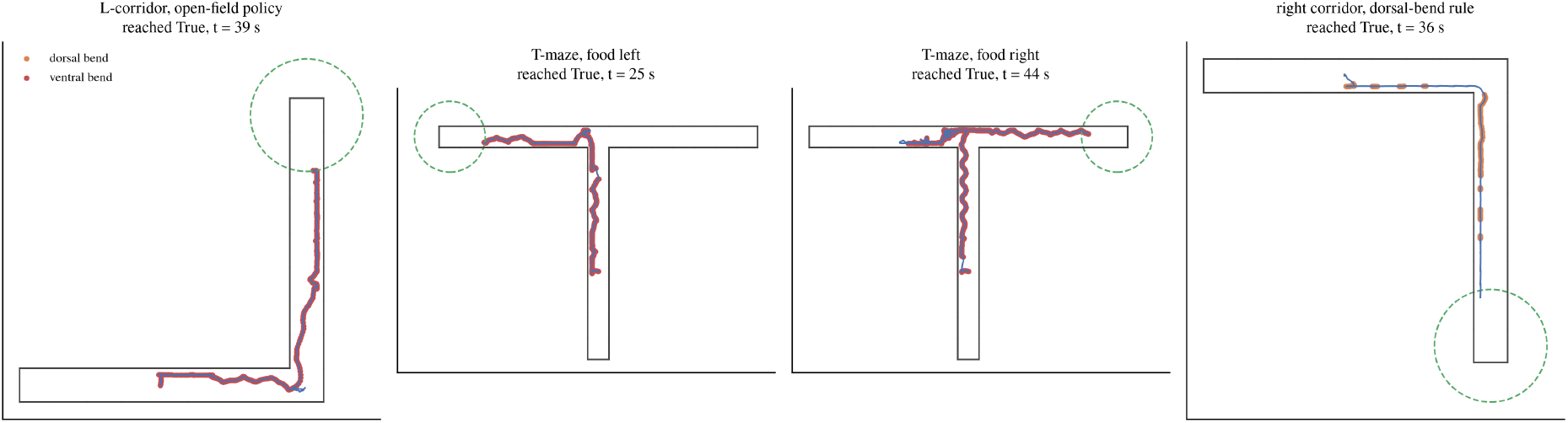
Mazes. Recorded trajectories of the open-ground policy in the L-corridor, of the maze-trained policy in the T-maze with food on the left and on the right, and of the right-turn-trained policy under the SMD-direction rule in the right-turn corridor. Dorsal and ventral deep-bend pulses are marked along the head path; the dashed circle is the goal radius.

The wall converts a blocked omega turn into a slide along the open side, so the dorsal or ventral direction of the bend is the direction of the turn (Fig 5). Under the ventral-only rule the open-ground policy passed left-turn corridors in 97–100% of jittered starts and right-turn corridors in 0–6% (0, 0 and 2 of 32 at widths 0.25, 0.30 and 0.35 mm). Policies with ventral-only bends entered the left (ventral) arm of a T-maze first in 127–128 of 128 jittered starts regardless of where the food was. A 3 *×* 3 grid maze that requires two right turns was not solved. Under the SMD-direction rule, which permits dorsal bends through whitelisted stimulation, retraining on the right-turn corridor succeeded (0.97–1.00 across widths) but produced a one-handed policy that failed the left-turn corridor (0.03–0.06), and policies not trained on right turns kept turning left. The rule preserved open-ground chemotaxis (0.871, CI 0.824–0.907). The retrained open-ground policy used dorsal bends 0.77 of the time and ventral bends 0.11, the opposite of real animals, in which 24% of sharp turns are dorsal [13]. The model has no dorsal–ventral cost asymmetry, so its preferred direction is set by initialization; the ventral preference of real worms lies outside what this model can explain.

**Figure 5:**
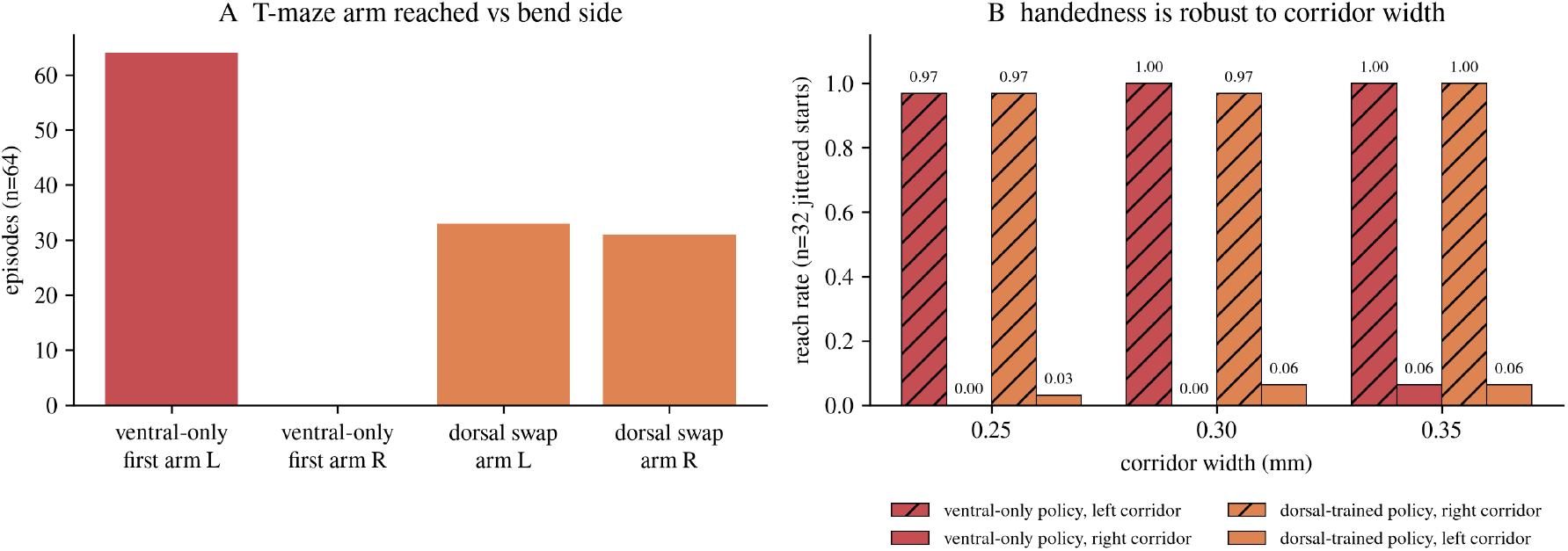
Bend direction is turn direction. (A) First arm reached in the T-maze from the fixed start by the open-ground policy with ventral-only bends and with the bend direction of the same policy swapped to dorsal (a control outside the whitelist); single-trajectory recordings over the two goal placements, not independent trials. (B) Reach in left- and right-turn corridors at widths 0.25–0.35 mm for the ventral-only policy and for the policy retrained on right turns under the SMD-direction rule (*n* = 32 validated jittered starts each).

### At a T-junction temporal sensing cannot choose the arm, so the policy enters one arm and reverses when wrong

Policies trained in the T-maze reached the food in 128 of 128 jittered starts under every rule and observation set, but they never chose the arm at the junction. First entry was into the left arm in 127–128 of 128 starts for every trained policy, and food on the right cost 5–10 s more than food on the left (for example 25.5 s versus 32.5 s for the first maze-trained policy). The correction was not a discrete choice but repeated 2 s reversal bouts that carried the worm tail-first back through the junction and into the other arm. Wall-contact observations were unnecessary. Zeroing them at evaluation left reach at 1.00 and first entry unchanged (median time 29.5 → 36.5 s); a policy trained with odor only also reached 1.00 (first entry left in 127 of 128, median 35 s). The open-ground policy without maze training reached 0.87 (right-hand goal 0.75, median 92.5 s against 32.5 s after maze training), so maze training mainly speeds up the correction, and a policy trained with contact only converged to stillness (0.00). In this model the “multisensory” requirement of real maze navigation [11] is met by odor plus wall physics, not by wall sensing.

The reason is informational. At the junction both arms present the same concentration to a nose that senses only the temporal history, so the correct arm can be found only by entering one. Adding proprioception (head curvature) raised open-ground reach to 0.957 and made T-maze navigation faster (median 20 s) but did not change first entry (128 of 128 left) and did not produce a weathervane (steering–source agreement 0.37); the policy used curvature to time the deep bend to the head-swing phase. The pre-registered prediction that dorsal bends would let the policy choose the arm (H5-6) was rejected: dorsal bends give the repertoire, not the decision. The stall reflex that takes corners also aborts a correctly aimed right entry. Recording body frames at the junction showed that a dorsal pulse carried the head into the right arm to *x* ≈ 0.45 mm, where it stalled against the wall. The stall registered as a weak fall in concentration (Δ log *C* of −0.13 to −0.15 per step), which triggered the open-ground reflex, reverse with a ventral head bias, and returned the worm to the left arm. Gating that reflex off during frontal wall contact reduced arm entry from 0.90 to 0.41 of jittered starts and reach from 0.48 to 0.30 for the lateral-information policy without maze training (introduced below), and in the fixed-start recordings masking SMDV at the junction abolished completion while masking AVA left only the left-hand goal reachable. The same reflex is thus the means of turning corners and the cause of the aborted entry.

### Lateral concentration information selects the bend direction

When the observation included the log-concentration difference between two points 0.1 mm to the left and right of the nose, an idealization of the head-swing sampling of real animals, the policy used it (Fig 6A). On open ground it reached the source in 98.8% of trials with a median of 16.5 s, the speed of the oracle. It used both bend directions (2,180 dorsal and 3,251 ventral pulses in 128 episodes), and the realized direction of the deep bend matched the side of the source in 0.73 of pulses, against 0.62 for the same rule without lateral information (chance 0.5). Because pulses within an episode are not independent, we recomputed the statistic per episode: 0.73 *±* 0.08 (all 128 episodes above 0.5) versus 0.61 *±* 0.18 (88 above, 35 below, 5 at 0.5), Mann–Whitney *p* = 2.6 *×* 10−8, Cohen”s *d* = 0.85. This reproduces, in the model, the sensory-guided choice of dorsal versus ventral turns reported for real animals [10]. In the T-maze, however, lateral information did not produce arm choice. Without maze training the lateral-information policy entered the right arm first in only 9 of 115 decided entries and reached a right-hand goal in 0.30 of starts; in the fixed-start recording it fired dorsal pulses toward a right-hand goal in 0.80 of cases, but each entry was aborted by the stall reflex described above. The maze-trained version abandoned dorsal pulses, entered left first in 128 of 128 starts, and instead reduced the cost of a right-hand goal to 5 s (22 versus 27 s).

**Figure 6:**
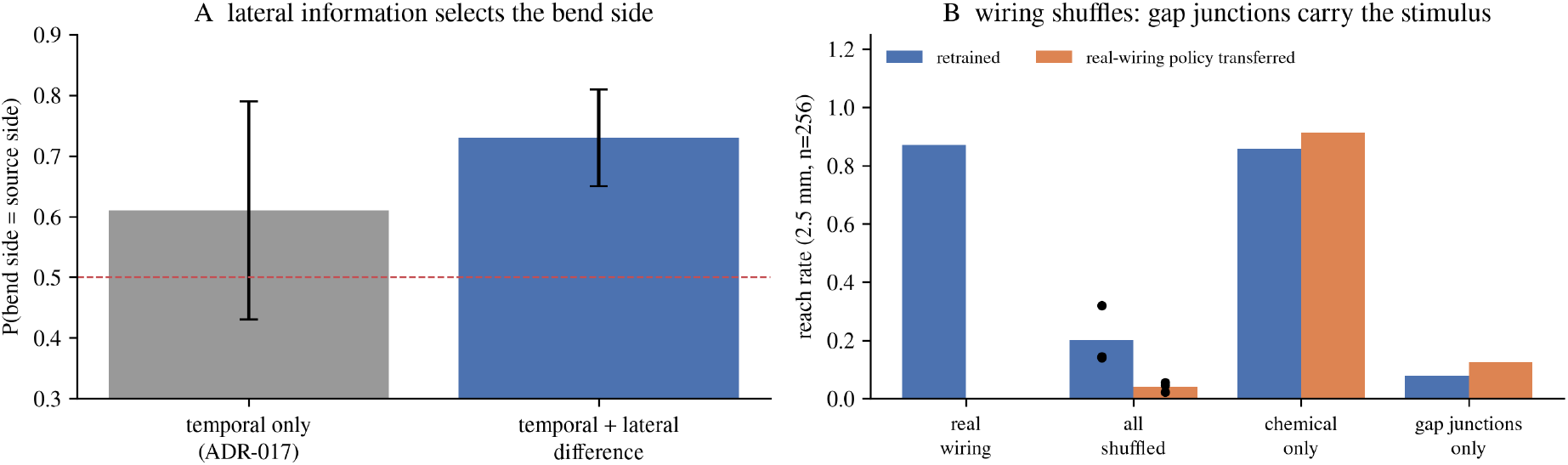
Lateral information and wiring controls. (A) Per-episode probability that the deep bend is toward the source, with and without lateral information (mean *±* s.d., 128 episodes). (B) Reach rates for real, fully shuffled, chemical-only shuffled and gap-junction-only shuffled wiring, retrained and with the real-wiring policy transferred.

### The real wiring outperforms shuffled wiring, through its gap junctions

The wiring contributed even though the command neurons were stimulated directly (Fig 6B). We shuffled the postsynaptic targets of all neuron-to-neuron chemical synapse entries (3,364 after sign splitting, Methods) and one end of all 1,084 gap junctions, preserving every weight, count and sign, and retrained under the same curriculum and rules. Three shuffled networks reached 0.141, 0.145 and 0.320 (mean 0.202) against 0.871 for the real wiring, and the real-wiring policy transferred to the shuffled networks reached 0.023–0.055. The deficit tracked the deep bend: in 128 evaluation episodes the real network fired 6,960 pulses, the three shuffled networks fewer than 40, 670 and 3,599, in the same order as their reach rates. Direct stimulation raises the potential of the stimulated neuron regardless of wiring, so the wiring”s contribution lies in how the stimulus spreads to the readout neurons SMD and RIV.

In the fitted dynamics that contribution was carried by the gap-junction layer. Shuffling only the chemical synapses left performance intact (retrained 0.859, transferred policy 0.914, 6,493 pulses), whereas shuffling only the gap junctions collapsed it (0.078 and 0.125, 116 pulses); the confidence intervals of the two layer-specific controls do not overlap (Table 2). The fit to the atlas had scaled chemical conductances down by more than an order of magnitude while leaving gap junctions near unity (Methods), so the neuron-to-neuron chemical layer is nearly silent in this model by construction. The defensible statement is therefore about the atlas-fitted dynamics: in them, the stimulus reaches SMD and RIV through the gap-junction network.

**Table 2:** Reach rates with Wilson 95% confidence intervals (*n* = 256, 2.5 mm unless stated). Rows with equal counts share an interval.

| Condition | Reach | 95% CI |
| --- | --- | --- |
| Full circuit (seed 0) | 0.914 | 0.873–0.943 |
| Full circuit (seeds 1–4) | 0.973 / 0.301 / 0.969 / 0.934 | 0.945–0.987 / 0.248–0.360 / 0.940–0.984 / 0.896–0.958 |
| Sensory channels only, 1.5 mm | 0.109 | 0.077–0.154 |
| Random policy, 1.5 mm / 2.5 mm | 0.125 / 0.043 | 0.090–0.171 / 0.024–0.075 |
| Real wiring, SMD-direction rule | 0.871 | 0.824–0.907 |
| All connections shuffled, retrained (3 seeds) | 0.141 / 0.145 / 0.320 | mean 0.202 |
| All shuffled, transferred policy (3 seeds) | 0.023 / 0.055 / 0.047 |  |
| Chemical synapses shuffled, retrained / transferred | 0.859 / 0.914 | 0.811–0.897 / 0.873–0.943 |
| Gap junctions shuffled, retrained / transferred | 0.078 / 0.125 | 0.051–0.118 / 0.090–0.171 |
| Lateral-information policy | 0.988 | 0.966–0.996 |
| AVB/AVA/RIV only, bend-always seed | 0.328 | 0.274–0.388 |
| Retrained without RIV | 0.191 | 0.148–0.244 |

### The reported numbers are robust to integration precision

All headline numbers held at the fine integration step required by our reporting rule. Headline values are those measured at the training step (0.25 ms, single precision) on 256 starts. On a common set of 128 starts, integrating at 0.05 ms in double precision gave reach rates of 0.914 (versus 0.938 at the coarse step) for the open-ground policy, 0.875 (versus 0.844) for the SMD-direction policy, and, for the maze-trained SMD-direction policy from the fixed start, the same outcome under both settings (reach and left first entry for both goal placements); only its maze transit time changed (median 26.5 → 33.5 s). Wilson 95% confidence intervals for the key comparisons are given in Table 2.

## Discussion

The central result is a partition. Inside the fitted connectome lie the sufficiency of five command and steering interneurons for chemotaxis, the necessity of reversal, deep bend and steering together, and a contribution of the wiring that is carried by gap junctions. Outside it lie the sensory-to-command step, which the fitted network does not perform, and the two ingredients that decide arm choice in a narrow maze, the dorsal–ventral direction of the deep bend and a lateral sample of the gradient. The model does not say that the connectome is unimportant for these. It says that in an atlas-fitted network with a literature motor layer they must be supplied from elsewhere, and it names what they are.

The learned strategy converged on the pirouette rule of real animals [8] without being told it, under the narrowest channel set (AVB, AVA, RIV) as well as the full one. Convergence was not sufficiency. A run-and-tumble policy with a fixed reorientation of about 170° cannot navigate, and every successful policy combined the rule with SMD steering that modulates the bend angle; real pirouettes likewise combine reversals with omega turns of variable angle [8, 14]. The convergence suggests that the rule is a property of the task and of the motor repertoire rather than of a particular circuit. The necessity of a deep bend follows from temporal sensing, as the lateral-information experiment showed: with a left–right sample the worm went straight to the source at the oracle”s speed, using the weathervane mechanism [9] in the form of bend direction rather than of continuous steering. Our failure to learn a weathervane from proprioception plus temporal sensing at a 0.5 s decision interval is consistent with the head-swing sampling of real animals being faster than our observations.

The wiring-shuffle controls answer a natural objection: if the agent stimulates command neurons directly, perhaps the wiring is irrelevant. It is not. Random wiring with identical weights, counts and signs performed at a fifth of the real wiring after retraining, and the loss tracked the loss of deep-bend pulses, that is, of the spread of stimulation to RIV and SMD. The layer dissociation, intact with shuffled chemical synapses and collapsed with shuffled gap junctions, has a clear caveat: the atlas fit had already reduced chemical conductances thirty-fold. We therefore claim a property of the fitted dynamics, not of the animal. It is, however, the same property that the atlas reported directly, that gap junctions and extrasynaptic signaling explain much of what anatomy alone does not [5], which suggests that the fit and the shuffle test probe the same structure. Whether the chemical layer becomes necessary under a different fit is a question the pipeline can answer; our own attempt with separate inhibitory operating points did not improve validation and was recorded as negative.

The maze results make three predictions for real animals. First, in T-mazes with corridors a few body widths wide, the arm entered should be predicted by the dorsal or ventral direction of the omega bend executed at the junction, which can be read from high-speed video. Second, conditions that suppress dorsal turns, whose spontaneous fraction is about 24% and falls to 16% during boundary escape [13], should bias animals toward the ventral arm and delay reaching a goal in the other arm; the wild-type absence of side bias (decision index 0.03 [11]) then requires that dorsal bends be used at the junction at a compensating rate. Third, stalling against a wall should trigger a pirouette, because in the model the same reflex that takes corners aborts a wrongly aimed entry. The model also reinterprets the mechanosensory requirement of maze navigation [11]. Wall-contact observations were unnecessary for our agent, so the failure of *mec-4* mutants may instead reflect the use of wall contact for propulsion [15] or for memory formation rather than for arm choice, a distinction that can be tested by separating the two.

The model has limits that bound these claims. The motor layer, steering and deep-bend rules are literature-based assumptions outside the connectome; each is anchored to a published mechanism [6, 10], but their parameters were chosen, not fitted. The body has no wall-pushing propulsion [15], so it crawls slowly in corridors, and it has no dorsal–ventral cost asymmetry, so it cannot explain the ventral preference of real turns. Sensory neurons are stimulated as currents, not through odor transduction, so the negative sensory result is a statement about the fitted network, not about real sensory neurons. Odour passes through walls. One learning algorithm and one small policy were used, and the 3 *×* 3 grid maze had no robust solution: a single individual traversed it from the fixed start, but 0 of 64 jittered in-corridor starts. One of five training seeds converged on a reversal-only local optimum; the sufficiency claim concerns the existence of solutions through the wiring, which four seeds and every ablation confirm, not the reliability of evolution strategies.

The approach treats a connectome as something to be interrogated by an adaptive experimenter rather than as a parameter set to be tuned. Its answers are of two kinds. Positive ones, such as the sufficiency of a command-interneuron interface, identify minimal stimulation targets for behavior. Negative ones, such as the silence of the sensory step or the near-absence of right turns with ventral bends, identify where the model, or the animal, must contain something the wiring diagram does not show.

## Methods

### Network model

We used the OpenWorm c302 framework [4] at parameter level C2, which represents each of the 302 neurons and 95 body-wall muscle cells as a single compartment with leak, slow and fast potassium and calcium conductances, graded chemical synapses and ohmic gap junctions. We re-implemented its equations in JAX so that hundreds of worms can be integrated in parallel on a GPU. The reference network was generated by c302 as NeuroML from the published connectome [1, 2] and never edited: 2,279 neuron-to-neuron chemical synapses (2,079 excitatory and 200 inhibitory in the c302 defaults), 552 neuron-to-muscle synapses and 1,084 gap-junction entries, with synaptic weight equal to the number of anatomical contacts. Voltage is integrated by backward Euler and gating, synaptic and calcium variables by exact exponential updates; against NEURON the re-implementation agrees to 0.001 mV RMS at a step of 0.05 ms. Exploration and training used a step of 0.25 ms in single precision; the headline numbers were re-checked at 0.05 ms in double precision (Results).

### Fitted dynamics (“Vfit”)

Two modifications, recorded as decisions ADR-010 and ADR-011, define the working variant. First, the sign of each neuron-to-neuron chemical synapse was assigned from receptor and transmitter expression in the CeNGEN atlas [16] with the method of Fenyves et al. [17]; synapses with conflicting predictions (1,085 of 2,279) were split into an excitatory and an inhibitory half, giving 3,364 synapse entries. Second, six global parameters were fitted to the signal-propagation atlas of Randi et al. [5]: the log conductance scales of excitatory, inhibitory and gap-junction connections and of the neuronal leak, and the half-activation voltage and slope of the graded synapse. The atlas protocol was simulated as a 2 pA current for 500 ms in each of 172 stimulus neurons, with the response defined as the mean depolarization over 2 s relative to an unstimulated control. The loss combined the negative Pearson correlation, a soft rank loss on the area under the receiver operating characteristic curve (AUROC) between significant and non-significant responder pairs, and a penalty on network-wide up-states; it was minimized by Adam (learning rate 0.1, 50 steps) on 32 stimulus neurons and validated on the remaining 140. The fitted values *θ* = [−3.44, −6.25, 0.13, 0.69, 1.48, 0.77] scale excitatory conductances by 0.032, inhibitory conductances by 0.002 and gap junctions by 1.14, double the leak, and shift the synaptic threshold to +1.5 mV with a slope of 10.8 mV. On the validation neurons the fitted model reached an AUROC of 0.646 (95% CI 0.617–0.671) and a Spearman correlation of 0.131 (0.103–0.153), against 0.546 and 0.075 for anatomy alone; the split-half reliability of the atlas bounds the attainable correlation at 0.140. Lesioning the model showed that this predictive power comes from gap-junction diffusion (gap junctions alone 0.613, chemical synapses alone 0.552). Of the 150 significant inhibitory responses in the atlas, 124 (83%) connect neuron pairs with no one- or two-step chemical or electrical path in the connectome and cannot be produced by any synaptic parameter setting. Under these dynamics, static stimulation of AVB or AVA differentiates B-type from A-type motor neurons, but stimulation of touch neurons does not differentiate the command state. Class-wise fits and separate inhibitory operating points overfit (validation correlation fell to ≈ 0) and were rejected.

### Body

The body is a two-dimensional overdamped viscoelastic rod of 24 segments and 1.0 mm length lying on its side, so that dorsoventral bending is in-plane. Stretch stiffness is 5 *×* 104 and bending stiffness 5 in units normalized to the tangential drag coefficient, with a normal-to-tangential drag ratio of 40 for crawling on agar; the rod is integrated at 10 *µ*s. With an imposed traveling wave the rod crawls at 0.24 mm/s. Muscle activation is mapped from the 24 rows of c302 muscle cells. Walls are line segments with a repulsive potential 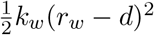 acting on any node closer than *r*_*w*_ = 60 *µ*m (*k*_*w*_ = 5 *×* 103); penetration was zero and results were stable for *k*_*w*_ between 2 *×* 103 and 2 *×* 104. Wall contact of the four head nodes within 0.12 mm is reported to the policy as three numbers (left, right, front) between 0 and 1.

### Motor layer and readouts

The c302 motor circuit has no intrinsic oscillator: no cell oscillates in isolation and no relaxation-oscillator regime exists in its channel-density space. The motor neurons and muscles were therefore replaced by the neuromechanical model of Boyle, Berri and Cohen [6] (ADR-012). It has twelve bistable units per side with on and off thresholds of 0.75 and 0.25, stretch-receptor gain 2.0 over 25% (forward) or 50% (backward) of the body, a muscle time constant of 100 ms and a maximal curvature of 6 mm−1, updated every 5 ms. Tuned in this way the body crawls at 0.18 mm/s at 0.50 Hz with head-to-tail propagation. Everything upstream of the command interneurons remains the fitted connectome. Four readouts couple the network to the motor layer, each anchored to a published mechanism. (i) Gates (ADR-012): the forward or backward gate opens when the mean depolarization of AVB or AVA relative to a 2 s unstimulated baseline exceeds 3 mV, the larger of the two winning. A quiescence gate for the sleep-active neuron RIS [12] stops the motor layer when RIS is depolarized by more than 3 mV and by more than 1.5 times any command or steering neuron. (ii) Head steering (ADR-013): the resting curvature of the anterior twelve rows, and the reference curvature of their stretch receptors, is offset by 0.03 mm−1 per mV of SMDD − SMDV depolarization, giving turns of about *±*50° in 12 s. (iii) Deep bend (ADR-016): when the RIV depolarization crosses 30 mV, a Gaussian curvature offset of width 4 rows and amplitude 0.36 mm−1 per mV above threshold travels from head to tail over 2.5 s; the pulse fires on the rising edge and re-arms only after RIV falls below threshold. The threshold separates direct RIV stimulation (≈ 55 mV) from gap-junction leak of SMD stimulation (≈ 20 mV). (iv) Direction (ADR-017): the pulse is dorsal when SMDD − SMDV exceeds 10 mV and ventral otherwise [10]. Network, motor layer and body exchange state every 50 ms.

### Environment, observations and reward

The odor is a two-dimensional Gaussian of peak 1 and width 2.0 mm centered on the source, evaluated at the nose; it passes through walls. On open ground the worm starts at the origin with a uniformly random heading and the source is placed at 1.5 or 2.5 mm in a uniformly random direction; in mazes each stage has one start pose, jittered as described below, and the T-maze goal arm is chosen at random. An episode lasts 40 s (120 s in corridors and the T-maze, 150 s in the grid) and ends when the nose comes within 0.5 mm of the source. Every 0.5 s the policy observes the log concentration level and its changes over the preceding 0.5, 1.0, 1.5 and 2.0 s, scaled by 10. Where stated it also receives the three wall-contact values, the mean curvature of the six head joints, or ten times the log-concentration difference between points 0.1 mm to the left and right of the nose. The reward per step is 100 Δ*C* − 0.05, plus 10 on reaching the source, minus 2 mm−1 times the final distance if the source is not reached. In the dwelling task the episode continues after arrival with +0.5 per step inside the radius and an optional cost of 5 per mm of head movement. Mazes are lattices of open cells of side equal to the corridor width (0.3 mm; 0.25 and 0.35 mm in the width test). Three stages were used: an L-shaped corridor with two 2.4 mm legs and one left or right corner; a T-maze with a 3.0 mm stem and 2.1 mm arms, with the source at the end of one arm chosen at random; and a 3 *×* 3 grid of 1.2 mm pitch. The width was chosen because the omega turn completes at 0.4 mm but is blocked at 0.3 mm.

### Policy, whitelist and training

The policy is a multilayer perceptron with one hidden layer of 16 tanh units and a sigmoid output per channel; the output scales a constant current between 0 and 6 pA injected into every neuron of the channel (both members of a bilateral pair) for the whole 0.5 s step. The whitelist (ADR-002) contains 19 channels and 35 neurons: sensory (PLM, ALM, AVM, ASH, ASE, AWC, AWA, ASK), command (AVB, AVA, PVC, AVD), steering (SMDD, SMDV, RMDD, RMDV, RIV) and state (RIS, ALA). Motor neurons and muscles are never stimulated. The command and steering set used throughout is AVB, AVA, SMDD, SMDV, RIV; the sensory set is ASE, AWC, AWA, ASK, ASH, PLM, ALM, AVM; the rule test used AVB, AVA, RIV; the dwelling task added RIS. Policies were trained with OpenAI evolution strategies [7]: antithetic Gaussian perturbations (*σ* = 0.2) of the mean parameter vector, fitness averaged over a fixed set of episodes shared by all candidates, centered rank normalization, and an update *θ* ← *θ* + (0.1*/σ*) *ϵF*. Stage 1 used 32 candidates *×* 8 episodes for 40 generations at 1.5 mm; stage 2 used 64 *×* 4 for 100 generations at 2.5 mm from the stage-1 policy (60 generations for the SMD-direction chain and the wiring controls). Maze policies started from the open-ground policy with zero weights on the new inputs and used 32 *×* 2 for 40 generations. The training seed was 0 for the main results and 1–4 for replication; evaluation always used 128 or 256 fresh episodes with seeds disjoint from training. Ablations mask a channel”s output at evaluation; sufficiency tests retrain without the channel. Maze evaluations deserve a caveat: the simulator is deterministic and each maze has a single start pose, so repeated episodes from that pose are identical (the T-maze differs only in the goal arm). All maze reach rates reported here therefore use start poses jittered within the corridor (along-axis *±*0.3 mm, lateral s.d. 0.03 mm, heading s.d. 3°), accepted only if all 25 body nodes lie inside the corridor; larger jitter placed the body outside the walls, through which odor passes, and was discarded. Corner-mechanism statistics and body-frame recordings are single-trajectory illustrations.

### Controls and statistics

The wiring shuffles permute the postsynaptic neuron of every neuron-to-neuron chemical synapse entry (3,364) and one end of every gap junction (1,084) with a fixed seed, preserving each connection”s weight, sign and kinetics, every presynaptic identity and out-degree, and all muscle connections; the layer-specific controls permute one class only. The oracle controller drives the motor layer directly with forward drive and a head bias proportional to the bearing to the source (gain 2, the best of the tested settings). A reversal onset is a step at which the AVA output crosses 0.5 upward; onset rates were computed within quintiles of the observed d*C/*d*t* over 128 episodes. Realised bend direction is the sign of the deep-bend pulse at the step it fires, compared with the side of the source. Reach rates are given with Wilson 95% confidence intervals; the per-episode bend-direction statistic uses a one-sided binomial sign test against 0.5 and a one-sided Mann–Whitney *U* test between policies. All hypotheses, criteria and verdicts were written down before the corresponding training run and are listed in Table 1 and in the repository.

## Code and data availability

Simulator, training and analysis code, decision records, logs and figures are available at https://github.com/kairess/worm-whisperer; recorded simulations with neuron read-outs can be viewed at https://kairess.github.io/worm-whisperer.

## Declaration of generative AI

A generative AI coding assistant (Claude, Anthropic) was used to write and run the simulation and analysis code and to refine the English of the manuscript. All experiments, results and text were reviewed and verified by the author, who takes full responsibility for the content.

